# Nuclear genome profiling of two species of *Epidendrum* (Orchidaceae): genome size, repeatome and ploidy

**DOI:** 10.1101/2025.09.17.675454

**Authors:** Miguel A. Alcalá-Gaxiola, Gerardo A. Salazar, Eric Hágsater, Mónica A. Flores-Iniestres, Lidia I. Cabrera, Aldo I. Aviña-Rivera, Pedro Mercado-Ruaro, Susana Magallon, Carolina Granados Mendoza, Aleida Núñez-Ruiz, Gloria Soldevila, Araxi Urrutia, Rubi N. Meza-Lázaro

## Abstract

Characterizing genomic properties such as genome size, ploidy level, heterozygosity, and repetitive DNA proportion and composition without relying on genome assembly is crucial for profiling the genomes of non-model species. Little is known about the nuclear genome of the large neotropical orchid genus *Epidendrum*. This study compares genome profiles of *Epidendrum anisatum* and *Epidendrum marmoratum,* using flow cytometry and k-mer analysis approaches, as well as bioinformatics ploidy level estimation and repeatome characterization. Multiple depths of coverage, *k* values, and k-mer-based tools for genome size estimation were explored and contrasted with cytometry genome size estimations. Cytometry and k-mer analyses yielded a consistently higher genome size for *E. anisatum* (mean 1C genome size = 2.59 Gb) than *E. marmoratum* (mean 1C genome size = 1.13 Gb), which represents a 2.3-fold genome size difference. Both species were identified as diploid with no evidence of strict partial endoreplication. The most important aspects to be taken into account to improve genome size estimation were heterozygosity, depth of coverage, and the maximum k-mer coverage. The genomes of both species were found to be highly repetitive (63–73%) and heavily dominated by Ty3-gypsy retrotransposons, particularly those of the Ogre family. Additionally, the genome of *E. anisatum* was characterized by the presence of a 172 bp satellite (*AniS1*), which represented 11% of the genome size. Together, both Ty3-gypsy transposons and *AniS1* shape the genome size difference between the two genomes. This study provides the first genome profiling for species in the genus *Epidendrum*, but also highlights the importance of using flow cytometry, cytogenetic approaches and bioinformatics techniques in combination for genome profiling.

## INTRODUCTION

Plant genomes exhibit a particularly large variation in their size, ploidy levels and repetitive DNA content (Wang *et al*. 2021). These characteristics play important roles in plant lineage evolution (Li *et al*. 2017) and are important considerations prior to whole genome sequencing and genome assembly efforts (Dominguez Del Angel *et al*. 2018). The characterization of genome size, heterozygosity, and genome repetitiveness, also referred to as genome profiling (Jenike *et al*. 2025), does not require genome assembly –a labor-intensive and complex process.

Genome size is usually estimated using laboratory techniques on living cells, specifically Feulgen densitometry and flow cytometry (Hardie *et al*. 2002; Leitch *et al*. 2019). These methods rely on staining techniques to quantify total nuclear DNA from live tissues. Flow cytometry is the gold standard for genome size estimation, but whole-genome endoreplication (also known as conventional endoreplication; CE) and partial endoreplication (PE) can challenge this method. CE and PE are cell cycle programs during which cells replicate their genomes without division (Brown *et al*. 2017; Shu *et al*. 2018). The first process results in somatic polyploid cells (Shu *et al*. 2018), and is widely distributed in plants, although some lineages are more prone to CE than others (Barow 2006; Trávníček *et al*. 2015; Pal’ová et al 2021). Differences in CE may also occur between individuals or even respond to environmental factors (Barow 2006). In contrast, PE results in cells that replicate only a fraction (P) of the genome (Brown *et al*. 2017) and it has only been reported in Orchidaceae (Brown *et al*. 2017). CE and PE can occur in one or several endoreplication rounds, and different plant tissues may have different proportions of 2C, 4C, 8C … nC or 2C, 4E, 8E, … nE nuclear populations, respectively. The 2C nuclear population sometimes constitutes only a small fraction in differentiated somatic tissues and can be overlooked by cytometry (Trávníček *et al*. 2015). Using plant tissues with a high proportion of the 2C population (such as orchid ovaries and pollinaria) can help overcome this difficulty (Trávníček *et al*. 2015; Brown *et al*. 2017).

Suitable tissues for cytometric analyses are often difficult to obtain, to preserve, or to process (e.g. Kim *et al*., 2024). Rapid bioinformatics methods, independent of whole genome assembly, are available for genome profiling, opening the possibility of exploring genome evolution in highly diverse non-model groups with no reference genome available, such as orchids, utilizing specimens from biological collections and museums. Bioinformatics techniques generally use genomic sequencing data to calculate the average depth of coverage in the dataset to estimate nuclear genome size, utilizing the principles of the Lander-Waterman equation (Lander and Waterman 1988; Pflug *et al*. 2020). These methods can be broadly divided into two types: read mapping to low-copy loci, which relies on a high-quality reference genome to identify low-copy loci (Pflug *et al*. 2020), and k-mer analysis, which is a reference genome-free technique for genome profiling.

K-mer analysis relies on plotting the number of times a substring of a fixed length k (k-mer) is found within the dataset (k-mer coverage), against the k-mer coverage frequency, to statistically estimate the genome size and other properties such as heterozygosity, sequencing error rate, and repetitive proportion. However, k-mer analysis is often distrusted for estimating nuclear genome size because of discrepancies with laboratory estimations (Hesse 2023). These discrepancies can result from several factors that can influence the final genome size estimation obtained from k-mer analysis. These factors can be methodological issues, such as the choice of the value of k (k value) during k-mer counting and the techniques used to estimate genome size from the resulting histogram. Additionally, there are intrinsic challenges related to the data and the genomic characteristics of the studied organisms, including high sequencing error rates, significant heterozygosity, the occurrence of polyploidy, and the amount of repetitive DNA in the genome (Pflug *et al*. 2020).

The abundance of repetitive DNA is the main factor behind the evolutionary changes in genome size across various eukaryotic lineages (Macas *et al*. 2015; Pellicer *et al*. 2018; Hloušková *et al*. 2019), alongside, to a lesser extent, polyploidization (e.g. Cordeiro *et al*. 2022). Some of the first methods for studying the repetitive DNA fraction of the genome relied on the reassociation kinetics of DNA using C_0_t curves (Britten and Kohne 1968). C_0_t curves are a biochemical approach that allows for the simultaneous estimation of genome size and the relative frequencies of repetitive DNA (Palmer and Black IV 1997). Massively parallel sequencing has introduced additional methods for DNA fraction estimation and repeat type identification. Repeat identification methods can be classified into *de novo* methods, methods relying on the comparison with sequence and structure databases, and hybrids of the previous two (Liao *et al*. 2023).

Repetitive DNA is typically classified into two types: tandem repeats, known as satellites, and interspersed repeats, referred to as transposons (Lee and Kim 2014). Transposons can then be classified into class I and class II transposons, based on whether an RNA intermediary is present or not, respectively. Long terminal repeat retrotransposons (LTRs) are the most common type of class I transposons in plants, particularly those belonging to the Ty1-copia and Ty3-gypsy superfamilies (Neumann *et al*. 2019).

The neotropical orchid genus *Epidendrum* L (subtribe Laeliinae Benth., subfamily Epidendroideae Lindl.) has previously been proposed as a promising model system for evolutionary and ecological studies (e.g. Pinheiro and Cozzolino 2013; Granados Mendoza *et al*. 2020; Pessoa *et al*. 2021), both because of its outstanding species diversity (over 1800 species; Karremans 2021; Moonlight et al. 2024) and morphological and ecological disparities (Hágsater and Soto 2005). Despite good morphological and ecological descriptions, little is known about their nuclear genome characteristics. This study aims to use two closely related endemic Mexican species, *Epidendrum anisatum* Lex and

*Epidendrum marmoratum* A. Rich. & Galeotti, to provide the first genomic profiling for this genus, focusing on the following aspects: 1) their genome sizes using both flow cytometry and bioinformatics methods; 2) different k-mer analysis approaches for estimating genome size; 3) bioinformatics ploidy level estimation and chromosome counting; and 4) the repeat composition, proportion, and abundance.

## METHODS

### Genome size estimation by flow cytometry

Leaf tissue samples of *E. anisatum* (voucher: *Salazar et al. 7370*) and *E. marmoratum* (*Salazar 10329*), and ovary tissue and pollinia (suitable for identifying the 2C and 1C nuclei population; Trávníček *et al*. 2015) were processed to estimate genome size by flow cytometry at the Laboratorio Nacional de Citometría de Flujo (LabNalCit; https://labnalcit.org/). The genome size was estimated following Doležel and Bartoš (2005). Nuclei were isolated in LB01 buffer (15 mM Tris, 2 mM EDTA, 20 mM NaCl, 80 mM KCl, 0.5 mM spermine tetrahydrochloride, 0.1% v/v Triton 300, and 15 mM β-mercaptoethanol, pH 7.5; Doležel 1989). The suspension was filtered through a 30–50 µm pore CellTrics™ filter (SYSMEX; https://www.sysmex.com) and treated with 50µg/mL RNAse A (Qiagen) and 50µg/mL propidium iodide (Sigma-Aldrich). Human peripheral blood mononuclear cells (PBMCs) from healthy individuals were used as a standard laboratory reference to calculate the *P. sativum* genome size. *Pisum sativum* and the *Epidendrum* samples were analyzed in a CytoFLEX S flow cytometer (Beckman-Coulter), individually and in combination with the internal references (PBMCs and *P. sativum*, respectively). Cytometry data analysis was performed using FlowJo® v. 10 (https://www.flowjo.com/). A genome size value for the *Epidendrum* samples was calculated as the average of the minimum and maximum 1C/2C values obtained from three replicates of the DNA content histograms of each tissue sample. Minimum and maximum values come from the interval of *P. sativum* estimations based on the human genome size range (human genome size range: 6.41-6.51; Piovesan et al. 2019).

### DNA sequencing

For library preparation, total genomic DNA was extracted from the leaf tissue of the same *Epidendrum* specimens noted earlier using the Qiagen DNeasy Plant Mini Kit. Twelve lysates of each sample were sequentially loaded onto a single DNeasy Mini spin column and centrifuged to obtain approximately 8 μg of DNA per sample. The quality and quantity of the genomic DNA were assessed by agarose gel electrophoresis and a NanoDrop 2000/2000c spectrophotometer (Thermo Fisher Scientific). DNA extractions were sent to Macrogen Inc. (Seoul, South Korea) for shotgun library preparation and sequencing. Libraries were prepared by random fragmentation and adapter ligation, and 150 bp paired-end sequencing was carried out in an Illumina HiSeq 2500 sequencer (Illumina, Inc., San Diego, California, US). This sequencing method produces suitable data sets without systematic biases, allowing the estimation of genome size and the proportion of repetitive DNA.

The sequences were trimmed and filtered using Trimmomatic 0.3.9 (Bolger *et al*. 2014) with the following parameters: ILLUMINACLIP:TruSeq3-PE-2.fa:2:30:10:2:True LEADING:3 TRAILING:3 MINLEN:100. We assessed the quality of the trimmed data using FastQC 0.11.9 (Andrews 2010).

### Genome size estimation by k-mer analysis

To assess the suitability of the whole dataset and estimate the minimum coverage required for genome size estimation, the depth of coverage of both datasets was calculated based on the flow cytometry 1C genome size values. Subsequently, the datasets were subsampled to simulate depths of coverage of 5×, 10×, 20×, 30×, and 40×. The depth of coverage of the datasets (both whole and subsampled) should be sufficient to resolve the error peak from the haploid peak (Ranallo-Benavidez 2020). Canonical k-mer counting and k-mer frequency plotting were performed using Jellyfish v.1.3 (Marçais and Kingsford 2011), with an exploratory maximum k-mer coverage value of 10,000 and subsequently a value of two million (Hesse 2023). To encompass a k-mer range other than the most common options (e.g. 21, 31), the k-values = 15, 17, 19, 21, 23, 25, 27, 29, 31, 33, 35, 41, 43, and 45 were used. Values are required to be odd to avoid palindromic k-mers. Genome size estimations were then carried out for both the whole datasets and each of the subsampled datasets.

Genome size for each k-value was estimated using three different software tools: CovEst 0.5.6 (Hozza *et al*. 2015), GenomeScope (Vurture *et al*. 2017), and findGSE 0.1.0 (Sun *et al*. 2018). Each uses a different statistical approach to calculate the depth of coverage, applying the principles of the Lander-Waterman equation. CovEst uses truncated Poisson distributions within a probabilistic framework, GenomeScope performs a non-linear minimum square optimization to adjust a mix of negative binomial distributions, and findGSE employs a mixed model, adjusting the k-mer frequencies iteratively using a skewed normal distribution. CovEst implements the “basic” and “repeats” algorithms, and findGSE implements the “hom” and “het” algorithms, which allow for examining the effects of high repetitive DNA proportion and heterozygosity, respectively. The mean absolute deviation (MAD) per tool with respect to the flow cytometry genome size estimation was calculated for each estimation using k-mer analysis. Heterozygosity was also estimated using GenomeScope and k = 21. Plots were generated using R 4.4.2 (R Core Team 2024) and the ggplot2 package (Wickham 2016).

### Ploidy level estimation

The ploidy level for each species was estimated using the distribution of bi-allele frequencies through nQuire (Weiß *et al*. 2018; https://github.com/clwgg/nQuire), which implements a Gaussian Mixture Model to test three possible ploidy models: diploidy, triploidy, and tetraploidy. To estimate the allele frequencies, the filtered sequences were mapped to the Angiosperm v.1 low-copy nuclear gene set of each species (Granados Mendoza *et al*. 2020) using BWA-MEM 2.2.1 (Li 2013) and SAMtools 1.15.1 (Danecek *et al*. 2021). The bi-allelic variants were filtered using the *denoise* command implemented in nQuire, and the ploidy level was assessed using the likelihood differences obtained from the *lrdmodel* command. Additionally, to confirm ploidy level, chromosome number was counted from developing root tips, which were pretreated using 0.002 M 8-hydroxyquinoline and fixed in absolute ethanol:glacial acetic acid (3:1). To prepare the slides, the root tips were hydrolyzed in 1 M HCl for 12 min at 60°, stained for one hour with Feulgen dye, and squashed carefully to disperse the cells. For the preparation of permanent slides, the liquid nitrogen technique was used; dispersed chromosomes were counted in intact cells.

### Repetitive DNA proportion and abundance estimation

The repeatomes of *E. anisatum* and *E. marmoratum* were characterized using the RepeatExplorer2 pipeline (Novák *et al*. 2020) within the graphical interface of the Galaxy platform (https://repeatexplorer-elixir.cerit-sc.cz/) and the default parameters. RepeatExplorer2 can be classified as a hybrid software for repeat detection (Liao *et al*. 2023), since it performs homology-based annotation, utilizing the RexDB databases (Neumann *et al*. 2019) and *de novo* detection through graph clustering (*seqclust* program) and TAREAN (Novák *et al*. 2017). TAREAN calculates the probability of a satellite using the C index (connected component index) and P index (pair completeness index), and its proportion based on the number of reads. The C index is a measure for identifying clusters representing tandem repeats (a value of one represents an exact tandem repeat). The P index is a measure of the proportion of unbroken read pairs (Novák *et al*. 2017).

Samples were analyzed using the individual and comparative protocols (Novák *et al*. 2020). The individual protocol allows for a detailed analysis of repetitive DNA and estimation of non-repetitive DNA (singlets) for a single species. The comparative protocol identifies shared clusters, allowing for comparisons by cluster instead of solely by repeat type. It does this by merging the reads from both samples, proceeding to the clustering and annotation steps, and finally separating the clusters by species. For the comparative protocol we used the same amount of reads for both species. The automatic annotation step was performed using the RexDB Viridiplantae 3.0 database, and the manual annotation step was based on the cluster’s graph shape analysis and information from the automatic annotation. The clusters corresponding to plastid or mitochondrial DNA were eliminated.

## RESULTS

### Genome size estimation by flow cytometry

Fluorescence intensity of DNA staining of the nuclei revealed a series of distinct peaks for *E. anisatum* and *E. marmoratum* derived from the ovary, pollinium, and leaf tissues (Fig. 1). Histograms showed multiple peak positions with no interpeak values, mirroring the profiles expected from a conventional endoreplication process. The pollinium histograms showed peaks corresponding to 1C and 2C, while the ovary tissue showed peaks at 2C and 4C. For *E. marmoratum*, the histograms indicated an additional 8C peak in ovary tissue and a 4C peak in pollinia (Fig. 1). The 1C value in gigabases (Gb; calculated from mass in pg) of *E. anisatum* ranged from 2.55 to 2.62 Gb (mean 1C value = 2.59 Gb) and that of *E. marmoratum* from 1.11 to 1.18 Gb (mean 1C value = 1.13 Gb; Supplementary Data Table S1).

**Fig 1.**
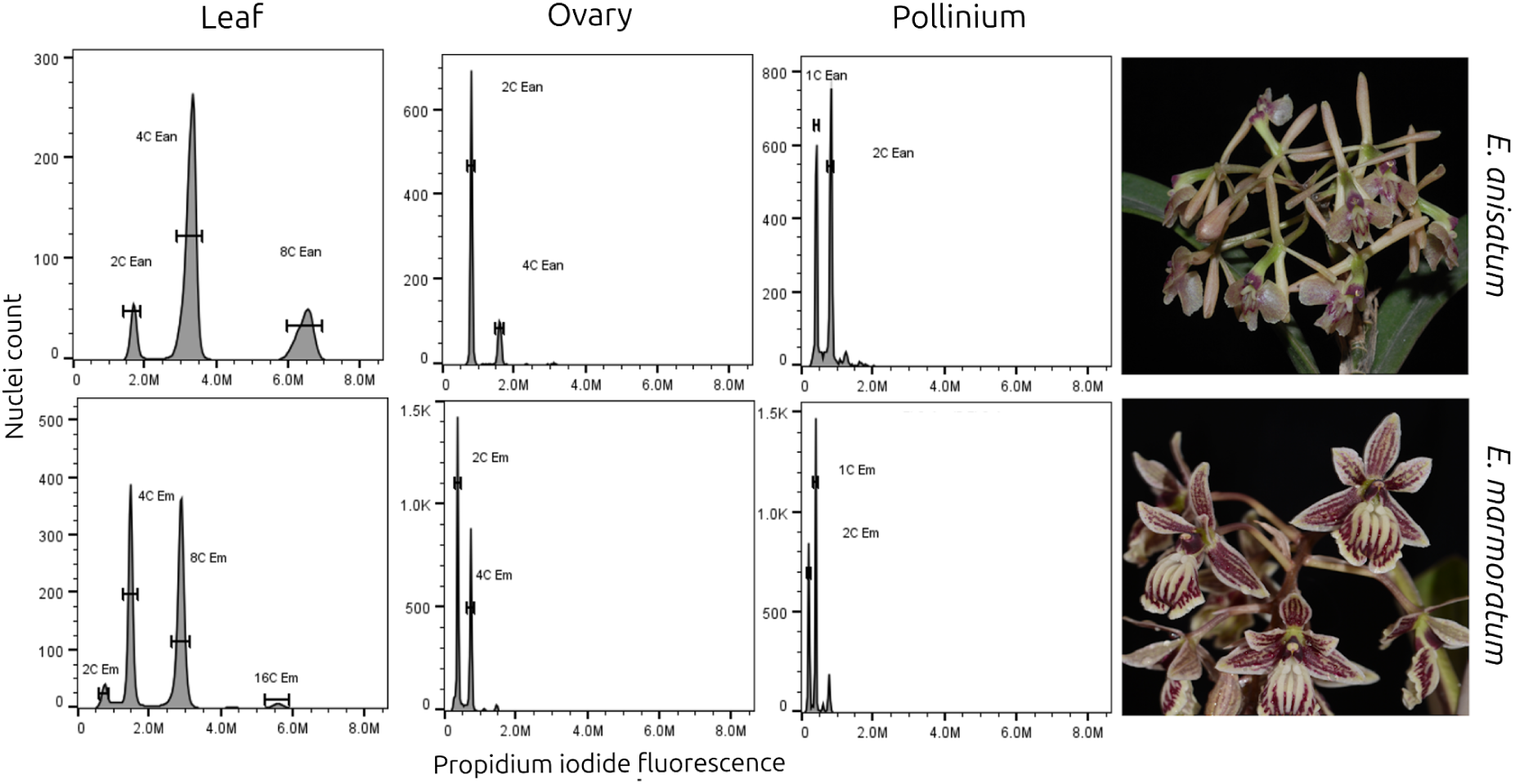
DNA content histograms for *E. anisatum* (voucher: *Salazar et al. 7370*; upper right) and *E. marmoratum* (voucher: *Salazar 10329*; lower right) from three different tissues: leaf (left), ovary (middle), and pollinium (right). Abbreviations: Ean, *E. anisatum*; Em, *E. marmoratum*.

### Genome size estimation by k-mer analysis and ploidy estimation

We obtained 760.0 million reads for *E. anisatum* and 765.3 million reads for *E. marmoratum* after trimming and filtering. The estimated maximum depth of coverage of the entire data set reached over 40× for *E. anisatum* and over 100× for *E. marmoratum.* The k-mer distribution histograms indicated that the error, heterozygous, and homozygous peaks can be clearly distinguished when using the whole datasets (Fig. 2). However, the error peaks were indistinguishable from the homozygous peak when analyzing low-depth of coverage datasets (5×). At higher coverage depths (greater than 10×), these two peaks became distinct, particularly in *E. marmoratum*. The homozygous and heterozygous peaks could be distinguished for some k-values at a 20× depth of coverage in *E. anisatum* (k-value = 17–21) and at a 30× depth of coverage in *E. marmoratum*. The error, the heterozygous, and the homozygous peaks could be clearly identified in the histograms for all k-values of *E. anisatum* starting at a 30× depth of coverage and over, and in the histograms of *E. marmoratum* at a 40× depth of coverage and over (Supplementary Data Figure S1).

**Fig 2.**
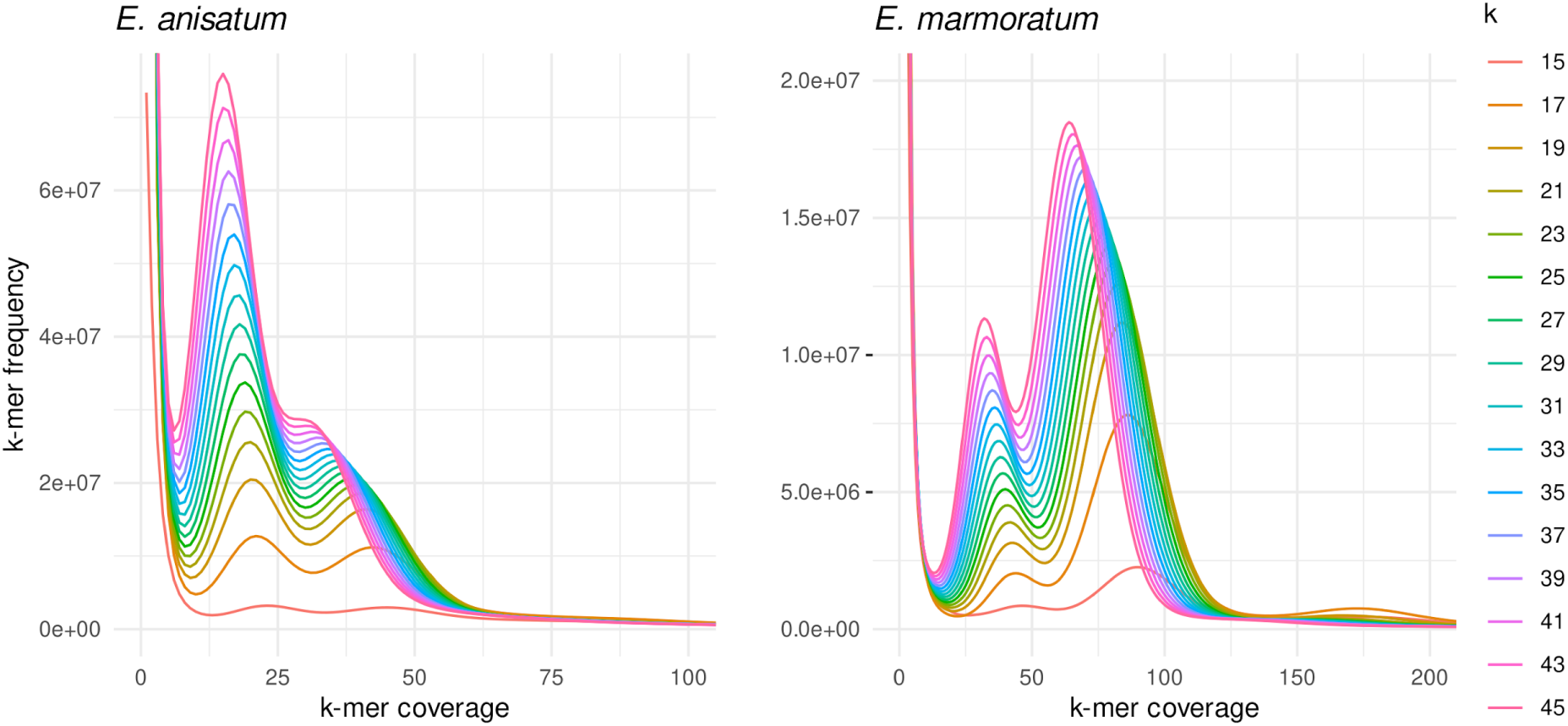
K-mer analysis histograms for *E. anisatum* (left) and *E. marmoratum* (right) for 16 values of k.

The full-sampling histograms (Fig. 2) illustrate the influence of the k-value on peak height and position. The heterozygous and homozygous peaks moved toward lower k-mer coverage (left), and the frequency of k-mers increases with higher k-values. In *E. anisatum,* the heterozygous peak was higher than the homozygous peak for every value of k (Fig. 2). This is mirrored by the heterozygosity estimation by GenomeScope, which was likewise higher in *E. anisatum* (1.7%) than in *E. marmoratum* (0.46%).

All of the tools estimated a higher genome size for *E. anisatum* than for *E. marmoratum*. The comparison of tools for genome size estimation yielded a general pattern in which genome size increases with the k-value (Fig. 3), mirroring the behavior of the homozygous peaks in the k-mer distribution histograms. The higher k-values tended to stabilize as the k-value increased. The lowest k-values (e.g. 15, 17) subestimated the genome size, which tended to be outliers with respect to the other k-values (Fig. 3) and, as a result, the genome size estimation interval is wider. The findGSE-het results (*E. anisatum*: 1.46–2.65 Gbp; *E. marmoratum*: 0.7–1.15 Gbp) had the lowest mean absolute deviation (MAD; 0.246 Gb for *E. anisatum* and 0.093 Gb for *E. marmoratum*; Fig. 3). In contrast, findGSE-hom substantially overestimated genome size for *E. anisatum* (2.85–5.3 Gbp; MAD = 2.04 Gb), while being more consistent with the flow cytometry estimation for *E. marmoratum* (0.8–1.3 Gbp; MAD = 0.100 Gb). GenomeScope was highly consistent with cytometry values, yielding the low MAD for *E. anisatum* (0.491 Gb) and *E. marmoratum* (0.101 Gb; Fig. 3). CovEst-basic tended to underestimate the genome size in both species (*E. anisatum*: 0.3–1.86 Gbp; *E. marmoratum*: 0.24–1.04 Gbp), and CovEst-repeats did not show a consistent increase in genome size with increasing k-values, instead exhibiting strong fluctuations (*E. anisatum*: 0.53–4.11 Gbp; *E. marmoratum*: 0.33–1.95 Gbp).

**Fig 3.**
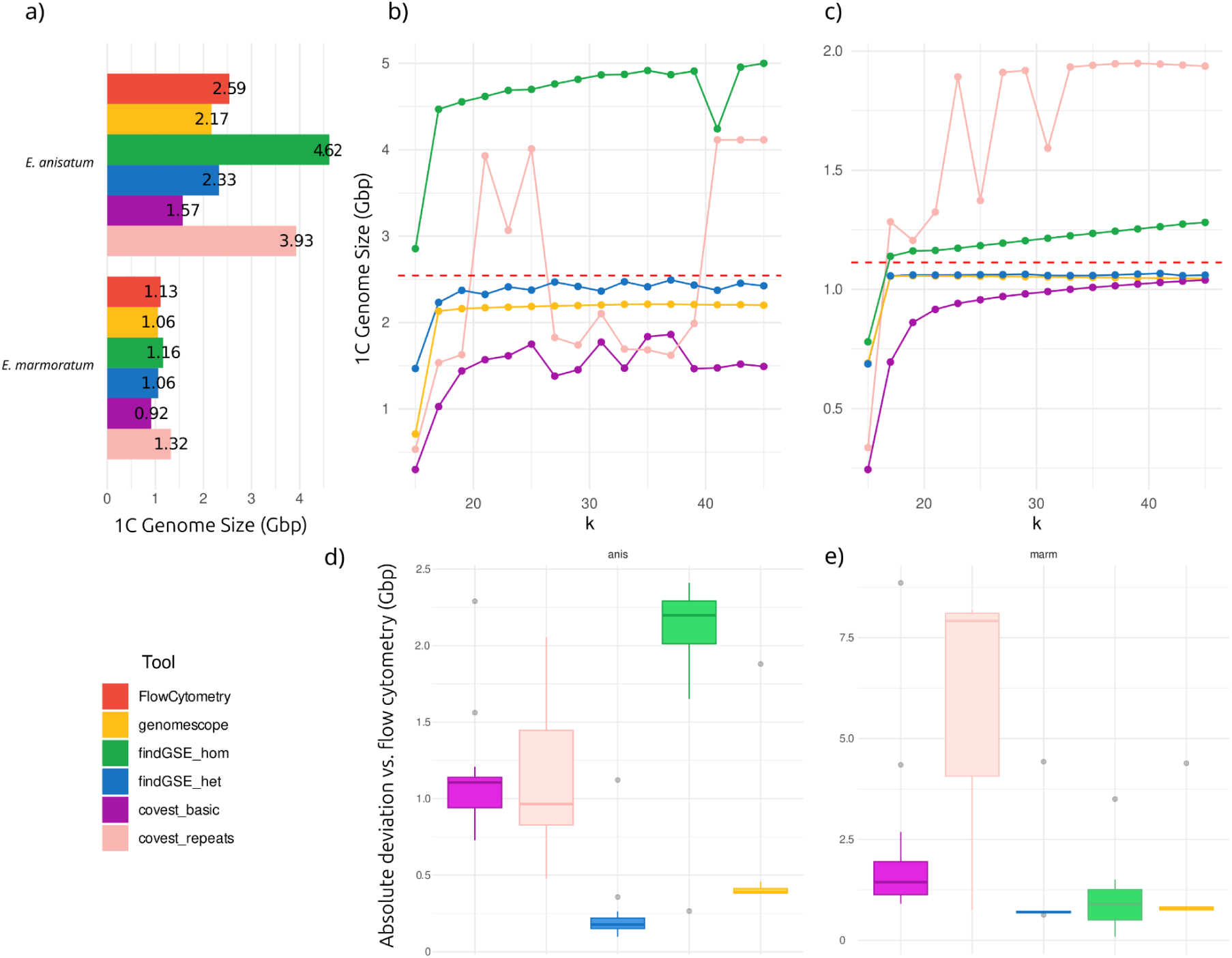
Genome size estimates for *E. anisatum* and *E. marmoratum* with flow cytometry and five bioinformatics tools. a) Comparison for k = 21, b) genome size estimates for *E. anisatum* with multiple values of k; c) genome size estimates for *E. marmoratum* with multiple values of k. The red dotted line represents the average flow cytometry value. Absolute deviation of genome size estimation per tool with respect to the flow cytometry estimation in d) *E.anisatum* and e) *E. marmoratum*.

The estimated genome sizes using a maximum count value of 10,000 (10K experiment) were generally lower for all tools in both species compared to using a maximum count value of 2 million (median of 2M experiment genome size - median of 10K experiment genome size= 0.24 Gb). The estimated genome size of the 2M experiment also tended to be closer to the flow cytometry genome size estimation with significantly lower MAD than the 10K experiment (Wilcoxon paired signed-rank test p = 0.0009). In the 10K experiment (Supplementary Data Figure S2; S3), the tool with the lowest MAD for *E. anisatum* was findGSE-het (0.546 Gb) and for *E. marmoratum* it was findGSE-hom (0.116 Gb).

The analysis of subsampled data simulating different depths of coverage yielded consistent genome size estimations similar to those of the whole datasets at depths of coverage ≥ 20× (Supplementary Data Figure S4). For both species, across most depths of coverage, findGSE-het was the most consistent tool with cytometry estimates, presenting the lowest MAD (Supplementary Data Figure S5). CovEst-basic and CovEst-repeats follow the same pattern as with the whole dataset estimations for both species in all samplings. CovEst-basic underestimates genome size, and CovEst-repeats yields results that strongly fluctuate with k value. The exception is for *E. marmoratum* at a 10× depth of coverage, where the CovEst-repeats estimations were always close to the genome size estimation of flow cytometry and had the lowest MAD of all the tools (Supplementary Data Figure S5). As the depth of coverage increased, the MAD tended to decrease for all tools, except for CovEst-basic and CovEst-repeats (Supplementary Data Figure S6).

We found both species to be diploids because the diploid model Δlog-likelihood was the lowest when compared with the free model likelihood (Table 1). Similarly, we counted 40 chromosomes for both species (Supplementary Data Figure S7).

**Table 1.** nQuire model comparisons.

| | Free model | $\Delta$ Diploid | $\Delta$ Triploid | $\Delta$ Tetraploid |
| --- | --- | --- | --- | --- |
| Species | log-likelihood | log-likelihood | log-likelihood | log-likelihood |
| *E. anisatum* | 9443.88 | **576.30** | 3668.26 | 4422.73 |
| *E. marmoratum* | 3556.02 | **644.05** | 1310.93 | 1422.86 |

### Estimation of repetitive DNA proportion and abundance

To estimate the proportion of repetitive DNA, the RepeatExplorer2 individual protocol determined a maximum number of analyzed reads (Nmax) corresponding to depths of coverage of 0.06× for *E. anisatum* and 0.43× for *E. marmoratum*. According to the results of this protocol, both species have highly repetitive genomes, with 73% and 63% repetitive content, respectively (Table 2). These proportions correspond to 1.72–2.04 Gb of repetitive DNA in *E. anisatum* and 0.62–0.78 Gb of repetitive DNA in *E. marmoratum*, according to the flow cytometry estimation (Fig. 4).

**Fig. 4.**
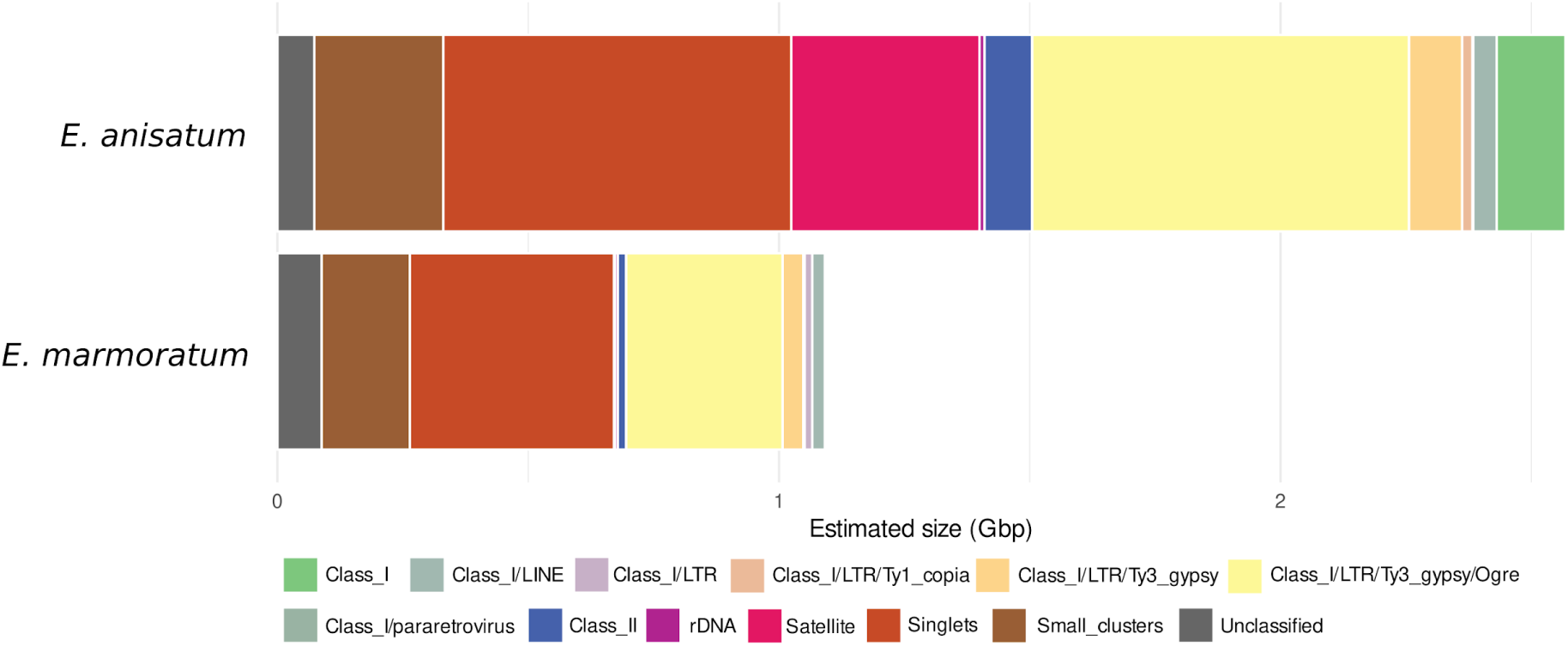
Abundance of repetitive DNA for the two species of *Epidendrum* analyzed after automatic annotation and manual review in RepeatExplorer2.

**Table 2.** Detailed repeat composition breakdown and comparison by genome proportion.

| Sequence type | Transposon classification | *E. anisatum* (%) | *E. marmoratum* (%) |
| --- | --- | --- | --- |
| **Singlets** |  | **27** | **37** |
| **Repeats** |  | **73** | **63** |
| Unclassified repeats |  | **2.85** | **7.99** |
| Transposons |  | **45.1** | **37.93** |
|  | **Class I** | **41.4** | **36.43** |
|  | Class I Unclassified | 5.33 | 0.43 |
|  | LINE | 1.83 | 2.28 |
|  | LTR | 34.24 | 33.72 |
|  | LTR Unclassified | 0.07 | 1.37 |
|  | Ty1-copia | 0.8 | 0.25 |
|  | Ty3-gypsy | 33.37 | 32.1 |
|  | Ty3-gypsy Unclassified | 0.58 | 0 |
|  | chromovirus/CRM | 1.3 | 2.13 |
|  | chromovirus/Reina | 0.24 | 0.001 |
|  | chromovirus/Tekay | 0.13 | 0.21 |
|  | non-chromovirus/Athila | 0.73 | 1.11 |
|  | non-chromovirus/Ogre | 29.24 | 28.34 |
|  | non-chromovirus/Retand | 1.15 | 0.31 |
|  | **Class II** | **3.7** | **1.5** |
|  | Subclass 1 | **3.7** | **1.5** |
|  | TIR Unclassified | 0.31 | 0 |
|  | EnSpm CACTA | 2.74 | 0.58 |
|  | hAT | 0.65 | 0.8 |
|  | MuDR Mutator | 0 | 0.11 |
| rDNA |  | **0.39** | **0.51** |
| Satellite |  | **14.6** | **0.19** |
| Small clusters |  | **10** | **16** |

\*Only showing repeat types with at least 0.1% genome proportion in at least one of the two species
\*\* After both automatic annotation and manual review

The dominant repeat type in both genomes was the Ty3-gypsy Ogre family, an LTR retrotransposon group that occupies 33.37% of the *E. anisatum* genome and 32.1% of the *E. marmoratum* genome. The most notable difference between the two species was in the abundance of a single satellite (henceforth *AniS1*), a 172 bp monomer with 44% GC composition that occupies more than 10% of the genome of *E. anisatum*, but is barely present in *E. marmoratum* (11.5% in *E. anisatum* and <0.01% in *E. marmoratum*). The satellite cluster has a C index (connected component index) score of 0.982 and a P index (pair completeness index) score of 0.485.

The genomes of the two species analyzed also differ in the proportions of unidentified repeats (2.85% in *E. anisatum* and 7.99% in *E. marmoratum*, differing in 5.14 percentage points) and unidentified Class I transposons (5.33% in *E. anisatum* and 0.43% in *E. marmoratum*, differing in 4.19 percentage points). Other repetitive DNA proportions show a difference of less than three percentage points between the two species.

The RepeatExplorer2 comparative protocol determined a max number of analyzed reads (Nmax) corresponding to depths of coverage of approximately 0.14× for *E. marmoratum* and 0.06× for *E. anisatum*. The results of this protocol showed a 67% total repeat proportion, which falls between the estimated repeat proportions of the two species according to the results of the individual protocol. However, the repeat proportions obtained by the comparative protocol for *E. anisatum* and *E. marmoratum* (76% and 57%, respectively) differed from those obtained through the individual protocol. The comparative protocol annotated 170 clusters, of which 19 were not shared between the two species: 17 were unique to *E. anisatum,* and two were unique to *E. marmoratum*. Additionally, in 16 of the shared clusters, one of the species dominated with at least 95% of the reads in the clusters. In 15 of these, *E. anisatum* dominated, while *E. marmoratum* only dominated in one. The results of the comparative protocol agreed with the results of the individual protocol in terms of the repeat composition of both species. The most abundant cluster was the same satellite cluster as in the individual protocol, which is shared but highly dominated by *E. anisatum*, accounting for more than 99% of the reads in the cluster. Similarly, the dominant repeat type in both species was the Ogre family of retrotransposons. A cluster-by-cluster comparison of the number of reads in both species using the comparative protocol showed that most Ogre clusters have similar proportions between the two species (Fig. 5).

**Fig 5.**
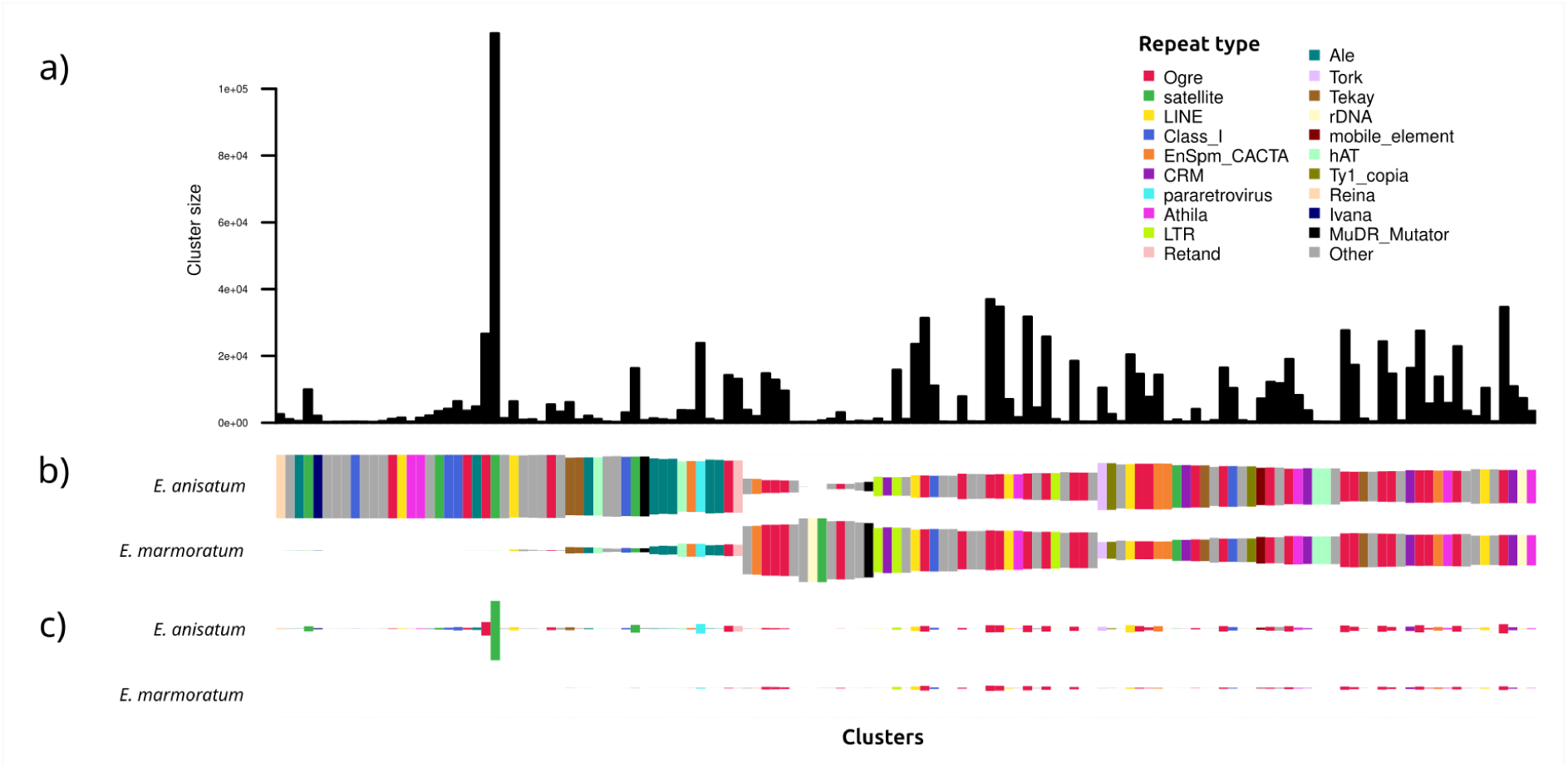
Results of the comparative protocol in RepeatExplorer2. a) Cluster size plot for all clusters, b) proportion comparison per cluster for *E. anisatum* and *E. marmoratum*, the rectangles represent what proportion of the cluster belongs to each species, c) abundance comparison per cluster for *E. anisatum* and *E. marmoratum*. In this case, the rectangles represent the absolute abundance of each cluster for each species.

## DISCUSSION

### Genome size estimation using flow cytometry

Genome endoreplication is a component of cell differentiation processes commonly observed in plants (Shu *et al*. 2018). In contrast, PE occurs mainly in orchids (Brown *et al*. 2017). Conventional and partial endoreplication can occur in multiple rounds, producing nuclear populations of 4C, 8C, 16C, and so on, and 4E, 8E, 16E, and so on, respectively (Brown *et al*. 2017; Loureiro *et al*. 2023). Endoreplicated nuclei can be more frequent than the 2C nuclei in some tissues (e.g., leaves and stems), which can be absent or undetected by the cytometer.

Ovary tissue and pollinia are less prone to endoreplication and are suitable for identifying the 2C nuclei population and estimating genome size (Trávníček *et al*. 2015). For example, although *E. anisatum* and *E. marmoratum* DNA content histograms have the same peaks in leaves and ovaries, the 2C peak is much lower than the 4C peak in the leaf histograms (Fig. 1). Such small peaks may be overlooked, and the genome size could be overestimated. The first peak in the pollinia histograms, corresponding to the reduced 1C gamete value, may cross-validate the genome size estimated from other tissues (Fig. 1). In *Epidendrum*, the 1C genome size estimation (*E. anisatum*: 2.41–2.81 Gb; *E. marmoratum* 1.11–1.29 Gb) resulted in half the 2C genome size calculated from both the ovary and leaf tissues (2.44–2.85 Gb; 1.05–1.22 Gb, respectively), suggesting that the first peak in the ovary and leaf was correctly identified and does not represent a round of endoreplication.

The ovary and leaf histograms did not present prominent intermediary peaks indicative of SPE, which agrees with a previous study in which no PE was found to occur in Laeliinae (Brown *et al*. 2017). In contrast, subtribe Pleurothallidinae Lind. ex G.Don, the sister group of Laeliinae, includes clades with evidence of PE (Brown *et al*. 2017; Chumová *et al*. 2021).

### Genome size estimation by k-mer analysis

Genome size estimation based on k-mer analysis is a powerful approach, but its accuracy can be affected by factors intrinsic to the genome, such as repetitive DNA proportion, heterozygosity, ploidy level, and genome size itself, as well as extrinsic factors associated with a particular dataset, such as sequencing error and dataset size. The choices of tools and parameters made during analysis can also deeply affect the genome size estimation and are closely related to the above mentioned intrinsic and extrinsic factors. The genomes of *E. anisatum* and *E. marmoratum* differ in terms of size, heterozygosity, and proportion of repetitive DNA, enabling us to assess how the choice of tools and their parameters influences the estimation of genome size, identify the main pitfalls, and improve the estimation.

A highly repetitive genome, such as that of *E. anisatum*, tends to have k-mers with high coverage values, representing the highly repeated parts of the genome. Selecting an appropriate maximum k-mer coverage value during the k-mer counting steps is the primary parameter for accurately estimating genome size. This parameter is genome-specific and tends to increase with the genome’s repetitive proportion (Hesse 2023). Although it underestimated genome sizes, the exploratory maximum k-mer coverage value of 10,000 provided a glimpse into the size, repetitiveness, and heterozygosity of the *Epidendrum* genomes analyzed. Increasing the maximum k-mer coverage to 2 million notably improved genome size estimation for both species. However, *E. anisatum* probably requires an even higher increase in this value to include the most highly repetitive k-mers in the histogram, as recommended by Hesse (2023).

To model repeats and sequencing errors in genome size estimation, CovEst provides the Model RE (Hozza *et al*. 2015) to obtain reasonable estimates of genome sizes from data at 1× genome coverage or less, using a k-value of 21. However, our k-value and simulated coverage using CovEst-repeats were only consistent and accurate at 10× depth of coverage for *E. marmoratum*, which may be evidence of the algorithm’s robustness when applied to real datasets with low depth of coverage and low heterozygosity.

The heterozygosity levels of the two species analyzed resulted in three distinct peaks; the first is the sequencing error peak, the second is the heterozygous peak, which occurs at exactly half the k-mer coverage value of the third peak, the homozygous peak (Hozza *et al*. 2015; Vurture *et al*. 2017; Sun *et al*. 2018). Distinguishing between error, heterozygous, and homozygous peaks is crucial for assessing the adequacy of coverage for genome estimation. The sequencing error peak was clearly observed in the histograms of *E. anisatum* at a depth of coverage of 20× or higher, and in *E. marmoratum* at a depth of coverage of 10× or higher, likely due to the smaller genome of *E. marmoratum*. However, a depth of coverage of 20×–25× or higher, generally recommended for diploid genomes (Vurture *et al*. 2017; Hesse 2023), performed better, allowing for the identification of heterozygous and homozygous peaks and enabling users to establish the parameters necessary for tools like findGSE-het to offset the impact of heterozygosity on the final genome size estimation. In contrast, when heterozygosity was not taken into account, findGSE-hom overestimated genome size, particularly in the highly heterozygous genome of *E. anisatum*.

The k-value of 21 is the most frequently suggested for genome size estimation (Hozza *et al*. 2015; Vurture *et al*. 2017; Pflug *et al*. 2020). Changing the k-value has been recommended for highly repetitive genomes (Vurture *et al*. 2017). This assertion is supported by the 10,000 maximum k-mer coverage experiment, which produces results closer to those of flow cytometry using higher k-values. In contrast, the two-million maximum k-mer coverage value estimations showed that k-values ≥ 17 produce very consistent genome size estimations with most tools (notable exceptions are Cov-Est repeats and findGSE-hom), emphasizing the importance of the maximum k-mer coverage value parameter.

K-mer analysis is a crucial tool for studying the genomes of non-model organisms that have not yet been fully assembled. For diploid organisms like *E. anisatum* and *E. marmoratum*, tools such as GenomeScope and findGSE-het, using k-values of 17 or higher, consistently provide estimates that align well with flow cytometry results. Learning about the complex genomic characteristics of a particular species through standard tools using default parameters can create a positive feedback loop that improves the choice of the tools and parameters for genome size estimation.

Besides ploidy level, heterozygosity, and the proportion of repetitive DNA, k-mer distribution can also be modified by endoreplication. Since endoreplication of the whole genome (CE) produces genome copies (as in preparation for cell division, but nuclear and cell division do not occur), we do not expect an effect on genome size estimates based on k-mer analyses. In contrast, PE alters coverage of a significant proportion of the genome, affecting k-mer distributions and genome size estimates (Piet *et al*., 2022). Species with PE might be challenging for k-mer-based methods of genome size estimation.

Genome size estimation for non-model species is considered a highly standardized approach. However, tissue availability and intrinsic genome characteristics (large genomes, polyploidy, endoreplication, and the proportion of repetitive DNA) can still preclude genome size estimation (e.g. Kim *et a*l. 2025) using cytometry and bioinformatic tools. Cross-validating flow cytometry and bioinformatics results might be particularly useful in those cases. For example, when only tissues suspected of showing significant conventional endoreplication, such as leaves, are available, bioinformatic tools can help to confirm that the first peak in cytometry histograms corresponds to 2C. Conversely, bioinformatic methods can be hindered by partial endoreplication, which flow cytometry can detect.

### Repetitive DNA composition in Epidendrum anisatum and E. marmoratum

*Epidendrum anisatum* and *E. marmoratum* have relatively small genomes, similar to those of most angiosperms, which typically have genomes smaller than 5 Gb/1C (Dodsworth *et al*. 2015). Repetitive DNA is considered the main driver of genome size variation in orchid nuclear genomes (Mehrotra and Goyal 2014), which span a 251-fold interval (from approximately 0.22 pg to 56.11 pg; Trávníček *et al.,* 2019; Chumová *et al*. 2021), but poliplodization can also be an important factor (*e.g. Epidendrum nocturnum* group; Cordeiro *et al*., 2022). Multiple cases of extensive chromosomal variation have been reported in *Epidendrum*, involving recent and ancient polyploidization, hybridization, dysploidy, and aneuploidy events (De Assis *et al*. 2013; Moraes *et al*. 2013; Cordeiro *et al*. 2022; Moraes et al. 2022; Nollet *et al*. 2022). However, ploidy and chromosome counts of both *E. anisatum* and *E. marmoratum* are the same (2*n* = 40) and are consistent with previous diploid counts reported for the genus (2*n* = 40; Felix and Guerra 2010). Therefore, it appears that repetitive DNA dynamics is the primary cause of the approximately 2.3-fold genome size difference between these two species. Class I/Ty3-gypsy retrotransposons significantly contributed to the approximately 600 Mbp size difference. However, the largest difference in proportion is in the abundance of the *AniS1* satellite, with a difference of more than 11 percentage points.

Repetitive DNA sequences are present in all higher plants and can account for up to 90% of the genome size in some species (Mehrotra and Goyal 2014). For example, previously reported repetitive DNA proportions in orchid genomes range from 9.94% in *Lepanthes clareae* (Chumová *et al*. 2021) to 88.9% in *Cymbidium goeringii* (Chung *et al*. 2022). The LTR transposon superfamilies Ty1-copia and Ty3-gypsy are the most abundant repetitive DNA types in a wide range of plant species (Neumann *et al*. 2019). Compared to other plant groups, LTR transposons are highly abundant in monocots, accounting for the majority of the genome (Kejnovsky *et al*. 2012) and have a very high turnover rate, with both gains and losses of transposable element (TE) sequences (*i.e.,* in grasses: Vicient *et al*. 2001; Giraud *et al*. 2021; in *Oryza*: Ma *et al*. 2004; Piegu *et al*. 2006; Vitte *et al*. 2007; in Pleurothallidinae orchids: Chumová *et al*. 2021).

In *E. anisatum* and *E. marmoratum*, Ty1-copia transposons are scarce (<1%) compared to other monocots (∼20% in *Zea mays* and ∼30% in *Oryza sativa*; Haberer *et al*. 2005). In contrast, Ty3-gypsy transposons are very abundant, resulting in a ratio of Ty3-gypsy/Ty1-copia of 41.7 in *E. anisatum* and 128.4 in *E. marmoratum*. Similarly, in the orchid subtribe Pleurothallidinae, the species with low Ty1-copia percentages have high Ty3-gypsy percentages (i.e. *Anathallis sanchezii* [Luer & Hirtz] Luer: Ty1-copia = 0.24%, Ty3-gypsy = 48.3%, 201.25 Ty3-gypsy/Ty1-copia; *Platystele misasiana* P.Ortiz: Ty1-copia = 0.02%, Ty3-gypsy = 59.46%, 2,963 Ty3-gypsy/Ty1-copia; Chumová *et al*. 2021). In contrast, some Poaceae groups have a Ty3-gypsy/Ty1 copia ratio which goes from 1.5 to 4 (*Avena* L.: Liu *et al*. 2019; *Lolium* L. and *Festuca* Tourn. ex L.: Zwyrtková *et al*. 2020; *Deschampsia* P.Beauv.: Amosova *et al*. 2022). This suggests that, in *Epidendrum* and Pleurothallidinae, Ty3-gypsy elements have higher rates of success in terms of amplification and subsequent retention in the genome than Ty1-copia elements (Kejnovsky *et al*. 2012).

Ty3-gypsy elements are frequently found in centromeric and pericentromeric regions, and may have an important structural role in heterochromatin (Jin *et al*. 2004; Neumann *et al*. 2011; Ma *et al*. 2023), particularly those with chromodomains in their structure (chromovirus, i.e. Tekay, CRM transposons; Neumann *et al*. 2011). Conversely, Ty1-copia elements tend to be more frequent in gene-rich regions (Wang *et al*. 2025A). However, Ty3-gypsy chromovirus elements can be found outside the heterochromatin regions (Neumann *et al*. 2011) and in *Pennisetum purpureum* Schumach. (Poaceae) Ty1-copia elements are more common in pericentromeric regions (Yu *et al*. 2022).

Most Ty3-gypsy elements in the two *Epidendrum* species belong to the Ogre family (87.6% in *E. anisatum*; 88.2% in *E. marmoratum*), a group of exceptionally large, plant-specific retrotransposons, which can reach up to 25 kb in length (Macas and Neumann 2007). Large expansions of Ogre elements have been documented in Fabaceae (*Lathyrus* L.: Ceccarelli *et al*., 2013; *P. sativum, Vicia faba* L.: Wang *et al*. 2025B), in Pleurothallidinae (e.g. 46% of the genome of *Anathallis sanchezii* are Ogre transposons; Chúmova *et al*. 2021) and in *Epidendrum* (Humberto *et al*. 2026). Although Ogre expansions are not well understood, it has been suggested that the expansion of certain transposons is related to host-directed epigenetic silencing of transposable elements (Huang *et al*. 2022) and the intrinsic characteristics of transposons (Wells and Feschotte, 2020).

The proportion of satellites marks the second most significant difference between *E. anisatum* and *E. marmoratum*. The repetitive fraction of *E. anisatum* is composed of approximately 20% satellite DNA. In a recent study, *E. anisatum* has been suggested to contain either 32.38% or 13% of its repetitive fraction as satellite DNA (Humberto *et al*. 2026, based on the RepeatExplorer2 individual or comparative protocol, respectively). The cluster corresponding to the *AniS1* satellite had a very good C index score (connected component index), whereas its P index (pair completeness index) score was relatively low, suggesting that *AniS1* is heavily dispersed throughout the *E. anisatum* genome. If this is the case, *AniS1* may be between the processes of dissemination and clustering (Ruiz-Ruano *et al*. 2016). Future fluorescence in situ hybridization (FISH) and third-generation sequencing studies will permit us to determine the distribution of *AniS1* across the *E. anisatum* genome.

Finally, another important difference between *E. anisatum* and *E. marmoratum* lies in the proportion of unclassified repeats (2.85% and 7.99%, respectively). This might have two different explanations: either no or not enough database hits could be found within the cluster; or there is conflicting evidence between different types of repeats (Novák *et al*., 2020). In the first scenario, it is likely that those clusters correspond to repeats which are very fragmented due to regulation by the host or decline due to being older elements. In the second scenario, the inability to classify the cluster is due to the dynamism of the repeatome. For example, satellites or rDNA can often be found interlaced with transposons (Lower *et al*. 2018; Šatović-Vukšić and Plohl 2021). For *E. anisatum* and *E. marmoratum*, the first scenario is more common, with only 4 unclassified clusters corresponding to the second case.

Ty3-gypsy retrotransposon and satellite proportions have been identified as the primary factors contributing to genome size variation in other plants. In Pleurothallidinae, the proportion of Ty3-gypsy retrotransposons seems to explain genome size variation (Chumová *et al*. 2021). However, in other plant groups, high percentages of satellite DNA have been found in species with large genome sizes (i.e. He *et al*. 2015; Pellicer *et al*. 2021). Future work should investigate to what extent LTR transposons and satellite DNA have been responsible for shaping genome size variation in different lineages of *Epidendrum*, analyzing a greater portion of its species diversity in an evolutionary context.

## CONCLUSIONS

Genomic profiling analysis of non-model species is an essential avenue for understanding the evolutionary history and genome dynamics of diverse lineages and is an essential step in high coverage whole genome sequencing and assembly of reference genome. This is particularly relevant in plant lineages where even related species may exhibit dramatic differences in genome size. In this study, we present the first analysis of the repetitive DNA composition in two species of the orchid genus *Epidendrum*, examining heterozygosity rates, ploidy and repeat content. We observed a 2.3-fold difference in genome size with no evidence of polyploidy but revealing a predominance of Ty3-gypsy Ogre retrotransposons and a unique satellite DNA, jointly responsible for the difference in genome size between the species analyzed.

This study also offers insights on the comparison of genome size estimations derived from flow cytometry and k-mer analysis, noting their respective strengths, consistency, and limitations in the context of the genomic characteristics of *Epidendrum*, including the apparent absence of strict partial endoreplication, heterozygosity variation, and a high percentage of repetitive DNA. Our findings underscore the importance of tailoring bioinformatic methods to the particular characteristics of each taxonomic group and establish a baseline for further research focused on a better understanding of the size and composition of the genomes of *Epidendrum* and their possible role in the diversification of this genus.

## Data Availability

All .histo files for genome size estimation, CLUSTER_TABLE.csv and SUPERCLUSTER_TABLE.csv files and Scripts are available at figshare: https://doi.org/10.6084/m9.figshare.c.8035945. Sequencing data analyzed for repeatome characterization is available at SRA (BioProject: PRJNA1330383).

## Acknowledgments

We thank Leonardo Pessoa Felix from the Centro de Ciências Agrárias, Universidade Federal da Paraíba, Brazil, for the *E. anisatum* micrographs and chromosome count; the Laboratorio de Biología Molecular, Instituto de Biología, Universidad Nacional Autónoma de México (UNAM) for access to DNA laboratory facilities; Eduardo A. Pérez García and the Orquidario Miguel Ángel Soto Arenas of the Facultad de Ciencias, UNAM, for providing material for cytometry analyses; and Cristian Cervantes, David Velázquez and Alfredo Wong from the Unidad de Sistemas y Tecnologías de la Información y Cómputo, Instituto de Biología, UNAM for support in using the HPC infrastructure. Access to the HPC cluster was made possible by the Instituto de Biología, UNAM, through Project Fronteras de la Ciencia 2016-01-1867, financed by Mexico’s Consejo Nacional de Ciencia y Tecnología (CONACYT). Computational resources for the RepeatExplorer analysis were provided by the ELIXIR-CZ project (LM2023055), part of the international ELIXIR infrastructure.

Supplementary Data Table S1. Flow cytometry genome size estimation statistics

Supplementary Data Fig. S1. K-mer analysis histograms for *E. anisatum* (upper row) and *E. marmoratum* (lower row) with 16 different values of k for five different depths of coverage sampling values (5×, 10×, 20×, 30×, and 40×).

Supplementary Data Fig. S2. Genome size estimates for *E. anisatum* and *E. marmoratum* using histograms generated from a k-mer counting with a maximum k-mer coverage value of 10,000 using 16 different values of k.

Supplementary Data Fig. S3. Absolute deviations in genome size estimates for *E. anisatum* (left) and *E. marmoratum* (right) per tool using the estimates generated from a k-mer counting with a maximum k-mer coverage value of 10,000 using 16 different values of k.

Supplementary Data Fig. S4. Genome size estimates for *E. anisatum* and *E. marmoratum* with flow cytometry and five bioinformatics tools using 16 different values of k and 5 different depth of coverage sampling values (5×, 10×, 20×, 30×, and 40×). An estimate of zero means that the program did not produce a result for that combination of values.

}

Supplementary Data Fig. S5. Absolute deviation boxplots for *E. anisatum* (upper row) and *E. marmoratum* (lower row) with 16 different values of k for five different depth of coverage sampling values (5×, 10×, 20×, 30×, and 40×).

Supplementary Data Fig. S6. Mean absolute deviation (MAD) changes as a function of depth of coverage for *E. anisatum* and *E. marmoratum*.

Supplementary Data Fig. S7. Chromosome count micrographs for *E. marmoratum* (left) and *E. anisatum* (right).

