## Supplementary Data for "Nuclear genome profiling of two species of *Epidendrum* (Orchidaceae): genome size, repeatome and ploidy"

Supplementary Data Table S1. Flow cytometry genome size estimation statistics

| Species | Plant tissue | 2C (pg) minimum (human 6.41 pg) | 2C (pg) maximum (human 6.51 pg) | CV sample | CV *P sativum* | 1C (Gb) minimum | 1C (Gb) maximum |
|---|---|---|---|---|---|---|---|
| *Epidendrum anisatum* | Polllinia | 5.27 | 5.36 | 3.36 | 3.36 | 2.58 | 2.62 |
| | Ovary | 5.22 | 5.30 | 2.92 | 2.92 | 2.55 | 2.59 |
| | Leaf | 5.27 | 5.36 | 3.70 | 3.80 | 2.58 | 2.62 |
| *Epidendrum marmoratum* | Polllinia | 2.38 | 2.41 | 2.87 | 2.87 | 1.16 | 1.18 |
| | Ovary | 2.24 | 2.28 | 3.15 | 3.15 | 1.10 | 1.11 |
| | Leaf | 2.23 | 2.27 | 5.09 | 4.06 | 1.09 | 1.11 |

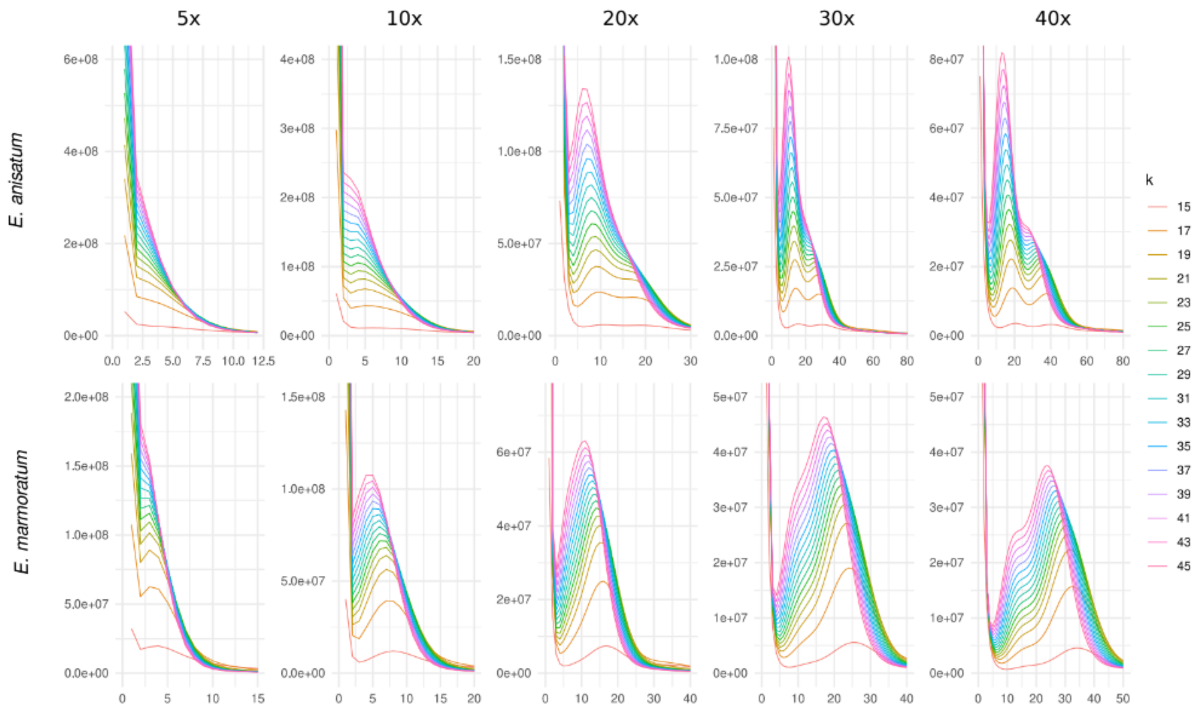


Supplementary Data Fig. S1. K-mer analysis histograms for *Epidendrum anisatum* (upper row) and *E. marmoratum* (lower row) with 16 different values of k for five different depths of coverage sampling values (5×, 10×, 20×, 30×, and 40×).

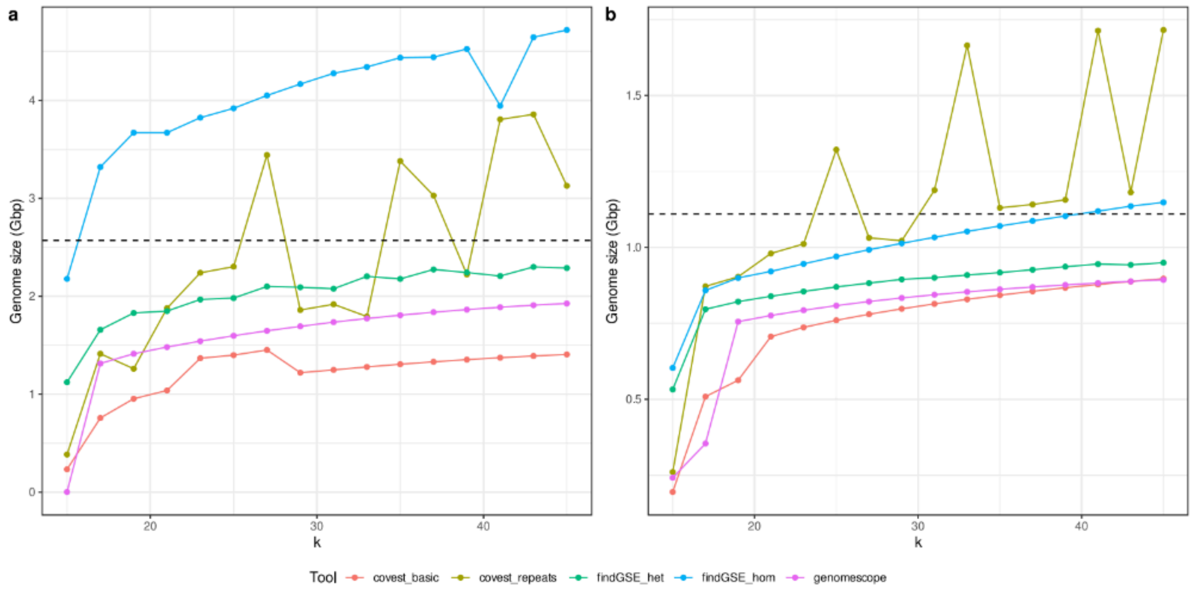


Supplementary Data Fig. S2. Genome size estimates for *Epidendrum anisatum* and *E. marmoratum* using histograms generated from a k-mer counting with a maximum k-mer coverage value of 10,000 using 16 different values of k.

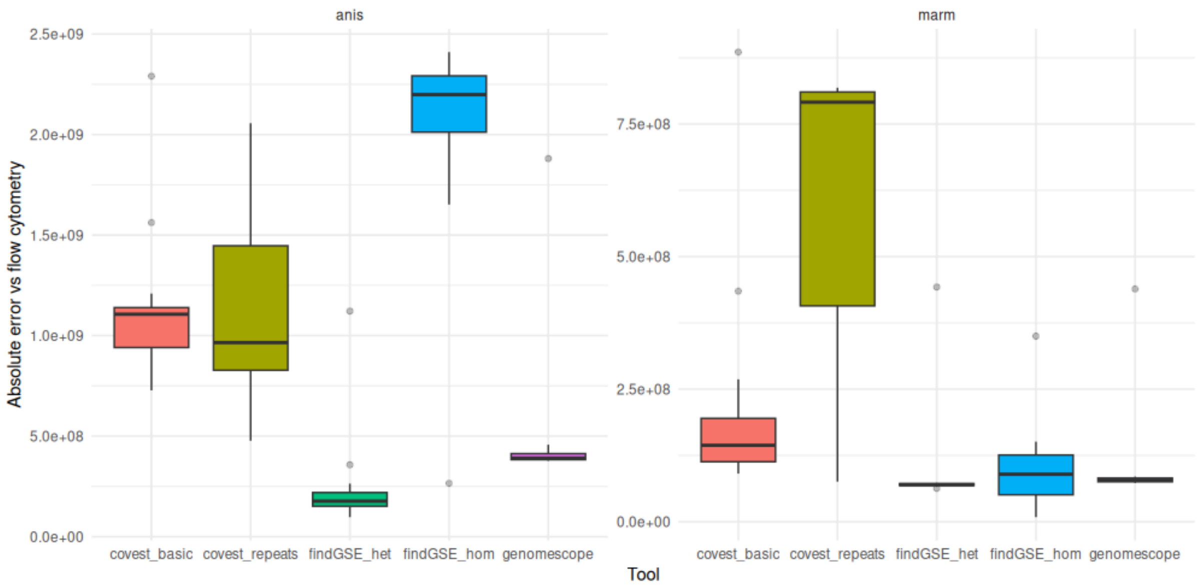


Supplementary Data Fig. S3. Absolute deviations in genome size estimates for *E. anisatum* (left) and *E. marmoratum* (right) per tool using the estimates generated from a k-mer counting with a maximum k-mer coverage value of 10,000 using 16 different values of k.

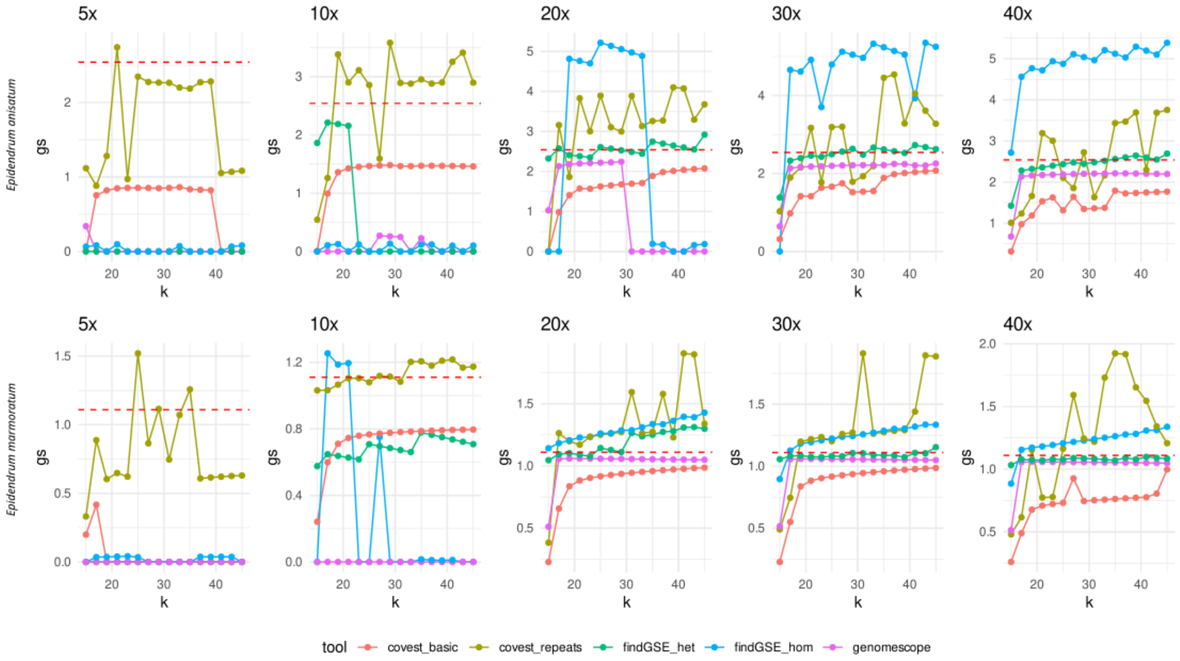


Supplementary Data Fig. S4. Genome size estimates for *Epidendrum anisatum* and *E. marmoratum* with flow cytometry and five bioinformatics tools using 16 different values of k and 5 different depth of coverage sampling values (5×, 10×, 20×, 30×, and 40×). An estimate of zero means that the program did not produce a result for that combination of values.

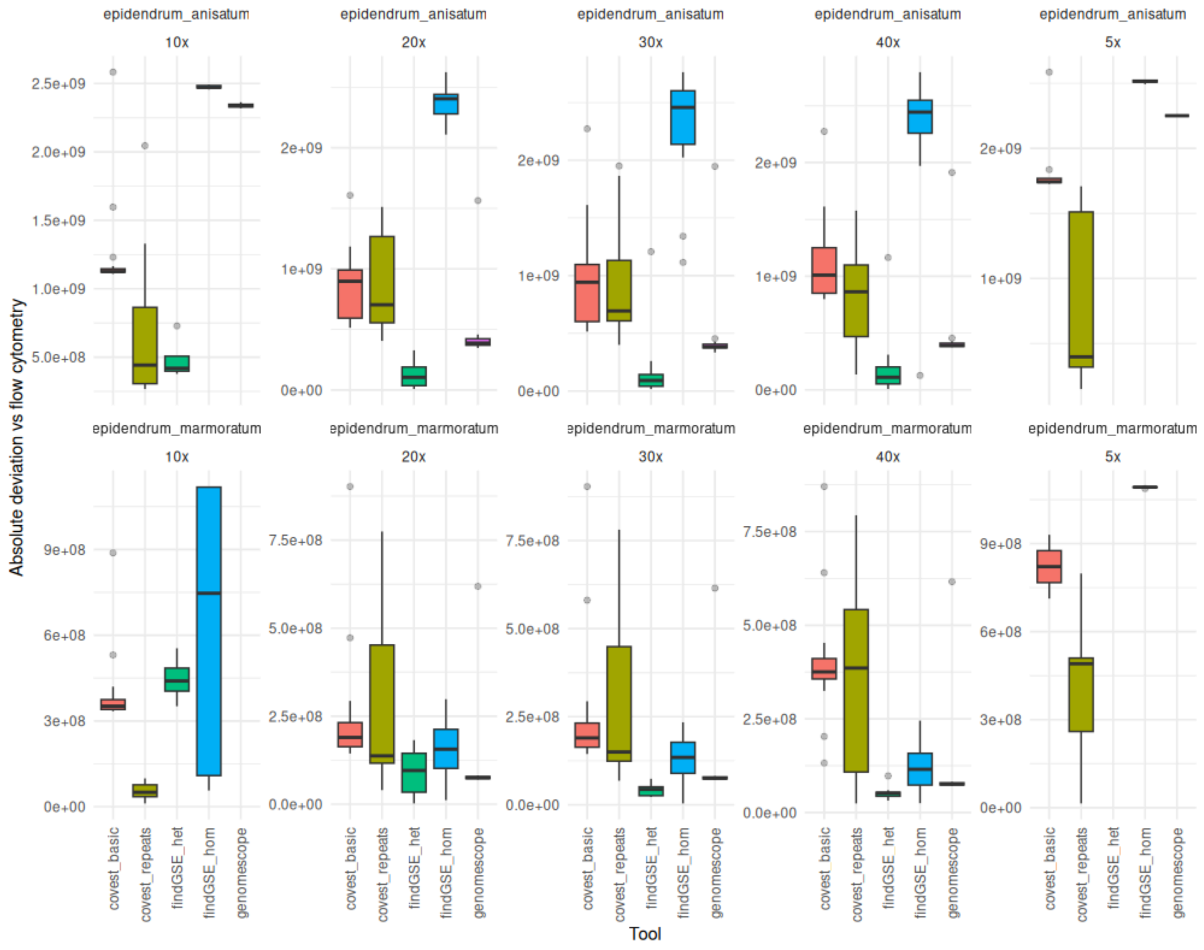


Supplementary Data Fig. S5. Absolute deviation boxplots for *E. anisatum* (upper row) and *E. marmoratum* (lower row) with 16 different values of k for five different depths of coverage sampling values (5×, 10×, 20×, 30×, and 40×).

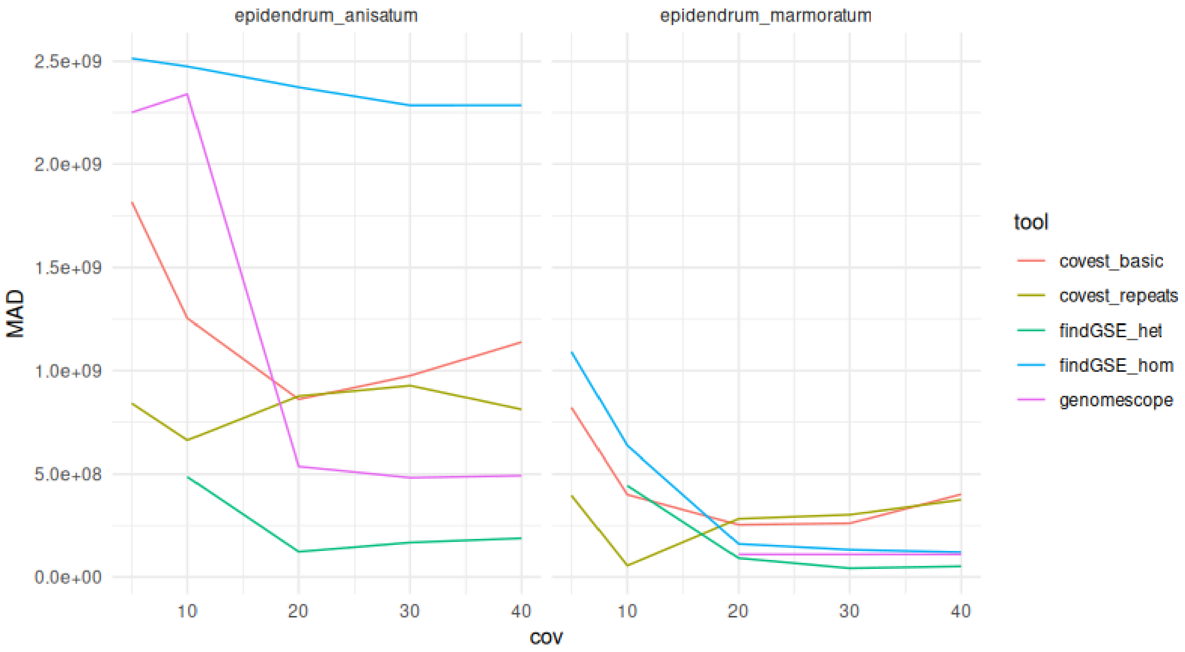


Supplementary Data Fig. S6. Mean absolute deviation (MAD) changes as a function of depth of coverage for *E. anisatum* and *E. marmoratum*.

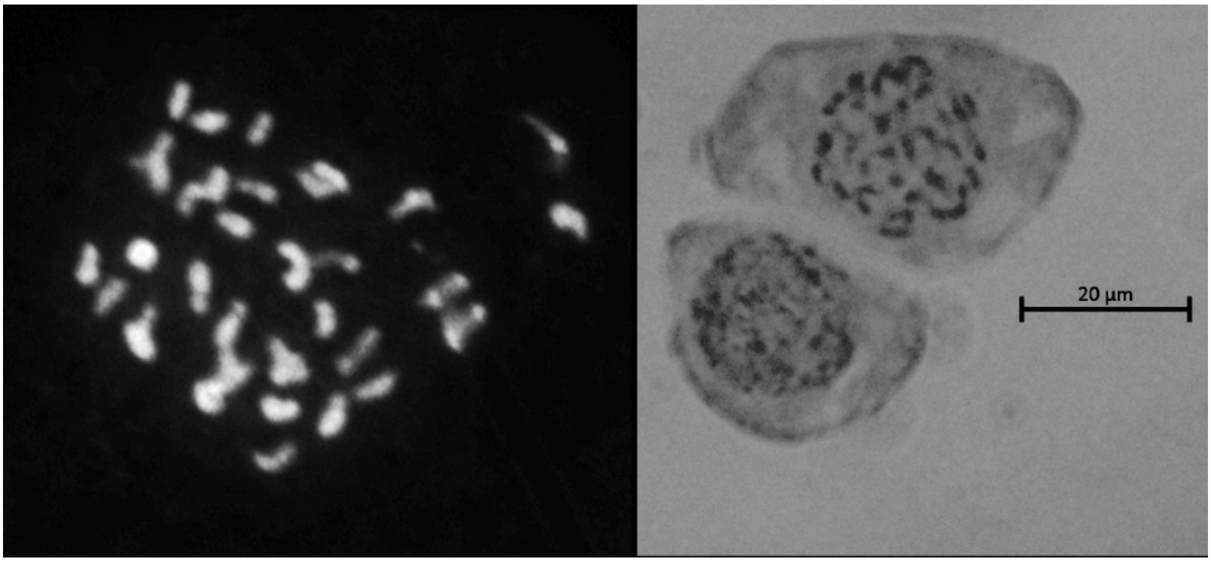


Supplementary Data Fig. S7. Chromosome count micrographs for *E. marmoratum* (left) and *E. anisatum* (right).
